# Regulation of the Na-K-2Cl cotransporter NKCC2 by ubiquitylation

**DOI:** 10.64898/2026.05.07.723577

**Authors:** Louise Nyrup Odgaard, Agnes Thoroee, Olivier Staub, Qi Wu, Robert A. Fenton, Lena K. Rosenbaek

## Abstract

NKCC2, localized to the apical membrane of thick ascending limb epithelial cells, is essential for renal salt handling and systemic electrolyte homeostasis. NKCC2 undergoes extensive ubiquitylation, with the E3 protein ligase Nedd4-2 implicated as a key regulator. However, progress has been limited by challenges expressing NKCC2 in mammalian cell lines, hindering mechanistic studies of NKCC2 ubiquitylation. Therefore, the aims of this study were to develop a mammalian cell model enabling mechanistic investigations of NKCC2 ubiquitylation, including the role of Nedd4-2 and the functional consequences of site-specific modification. A tetracycline-inducible MDCKI cell line was generated expressing human NKCC2 and used to assess Nedd4-2-dependent and site-specific ubiquitylation of NKCC2 using biochemical, imaging, and functional assays. The MDCKI cell line demonstrated stable, inducible expression of full-length human NKCC2. In this cell line, mutating the ubiquitylation site at K871 increased membrane abundance and uptake activity, without altering internalization rates. Nedd4-2 co-immunoprecipitated with NKCC2, and Nedd4-2 deletion increased total, but not membrane NKCC2 abundance. In summary, ubiquitylation on NKCC2 at K871 plays a key role in controlling NKCC2 membrane localization and thus function. Although Nedd4-2 can modulate NKCC2 abundance, it is not involved in NKCC2 trafficking. We conclude that the generated cell line provides a robust model for mechanistic studies of NKCC2 and will aid studies examining posttranslational regulation of NKCC2.

## Introduction

The sodium-potassium-chloride cotransporter 2 (NKCC2, encoded by *Slc12a1*), is a member of the *Slc12* family of Na^+^-dependent Cation-Chloride Cotransporters (CCC), which also includes the sodium-chloride-cotransporter, NCC and the sodium-potassium-chloride cotransporter 1, NKCC1 (1, 2). NKCC2 is expressed in the thick ascending limb of Henle’s loop (TAHL) where it is responsible for reabsorption of up to 25% of filtered Na^+^, K^+^, and Cl^-^ and hence plays a role in Na^+^/K^+^ balance and regulation of blood pressure (3). NKCC2 is also essential for generating the kidney’s corticomedullary osmotic gradient and urine concentration (4). Due to its role in generating a positive luminal potential, NKCC2 is important for Ca^2+^ and Mg^2+^ reabsorption by the TAHL. Furthermore, NKCC2 is expressed in macula densa (MD) cells, where it plays a central role in how MD cells sense and regulate glomerular filtration, known as tubuloglomerular feedback (5, 6). Supporting these roles, mutations in *Slc12a1* result in Type I Bartter syndrome, characterized by polyuria, hyponatremia, hypokalemic alkalosis, hypomagnesemia, hypercalciuria, and hypotension (6–11). NKCC2 is also the target of loop diuretics such as furosemide and bumetanide, which cause diuresis, natriuresis, kaliuresis and hypomagnesemia (8, 12, 13).

Under basal conditions, NKCC2 is localized both to the apical plasma membrane (14, 15), large subapical reservoirs and in the Golgi apparatus (5). Phosphorylation of NKCC2, through the With-No-Lysine-STE20/SPS1-related proline/alanine-rich kinase/Oxidative stress-responsive kinase 1 (WNK-SPAK/OSR1) signaling cascade (16), alters NKCC2 membrane abundance and is critical for its maximal transport capacity (17, 18). NKCC2 phosphorylation can also occur through protein kinase A (PKA) and AMP-activated protein kinase (AMPK) cascades (18–20).

Recently, evidence for NKCC2 ubiquitylation has emerged (21). NKCC2 contains a PY-motif (22), co-immunoprecipitates with the E3 protein ligase Nedd4-2 and NKCC2 ubiquitylation is increased in mice in response to high-salt diet (23). Ubiquitylation of NKCC2 is also enhanced by cGMP, mediated by the CRL (Cullin-RING-E3 ubiquitin Ligase) complex (24), resulting in proteasomal NKCC2 degradation (21, 25). Given that other *Slc12* family members are found to be ubiquitylated (26–29) and that ubiquitylation is involved in regulation of trafficking and surface stability of NKCC1 (30, 31) and NCC (32–34), similar mechanisms may apply to NKCC2. However, despite studies in TAHL suspensions with native NKCC2 expression having advanced our knowledge of NKCC2 biology (35–37), studies on the role of individual regulatory sites on NKCC2 have been hampered by the lack of suitable cell models. For example, to our knowledge, no investigations have succeeded in obtaining stable heterologous expression of functional NKCC2 in mammalian cells (3, 38–40) and transient transfection studies make comparisons of different mutants difficult.

Therefore, the initial aim of this study was to generate a polarized mammalian cell line with stable expression of full-length NKCC2 that responds to external stimuli with regulated NKCC2 trafficking and allows studies of individual sites within NKCC2. The second aim was to investigate the regulation of NKCC2 by site-specific ubiquitylation. Finally, we further examined the involvement of Nedd4-2 in NKCC2 ubiquitylation.

## Methods

### Generation of tetracycline inducible human NKCC2 MDCKI cell lines

Generation of the MDCKI cell line with tetracycline inducible hNKCC2A was performed as described (41). Briefly, the pcDNA5/FRT/TO/TOPO-hNKCC2 cDNA vectors were co-transfected with pOG44 (encoding flp recombinase) using Lipofectamine 2000 (Thermo Scientific) into a tetracycline inducible MDCK type I cell line containing a single FRT site in its genome (42). Positive clones were selected using 500 μg/ml Hygromycin B and the stable MDCKI-hNKCC2A cell lines were maintained in DMEM high glucose with 10% DBS, 150 μg/ml Hygromycin B, and 5 μg/ml Blasticidin HCl (Thermo Scientific). Unless otherwise indicated, cells were induced with 10 μg/ml tetracycline HCl and 5 mM valproic acid for 24 h at 37 °C prior to experiments. K-to-R mutations of ubiquitylation sites were introduced into human NKCC2 (isoform A, hNKCC2A) by site directed mutagenesis (Mutagenex).

### Immunoprecipitation of total cell lysates

Cells were grown in T25 flasks until confluent and induced. Cells were washed twice in ice-cold PBS-CM and subsequently lysed in IP lysis buffer (20 mM Tris-Base, 135 mM NaCl, 1% NP-40, 5 mM EDTA, pH 7.4) containing phosphatase and protease inhibitors, 22 μM PR619 (Abcam) and 20 mM N-ethylmaleimide for 30 min at 4 °C with mild agitation. Cell lysates were sonicated and centrifuged at 10,000 x g for 5 min at 4 °C. One fraction of the supernatant was retained for input sample. FLAG-M2 affinity beads (A2220, Merck) were washed in IP lysis buffer with 0.1% BSA for 1 hour at RT with rotation, before equal volumes of the remaining sample supernatants were added and incubated at 4 °C with rotation overnight. Beads were washed extensively in IP lysis buffer with inhibitors and immunoprecipitated proteins eluted using FLAG-peptide (DYKDDDDK, Genscript, 200 µg/ml) in TBS buffer (10 mM Tris-HCl, 150 mM NaCl, pH 7.4). For immunoblotting Laemmli sample buffer was added for a final DTT concentration of 15 mg/ml and samples were subsequently heated for 15 min at 60°C.

#### Mass spectrometry analysis

Immunoprecipitated NKCC2 proteins were subjected to filter-aided sample preparation and peptides KGG-motif enriched prior to LC-MS/MS and data analysis as described previously (43).

#### Nedd4-2 Knockout (KO) cells

Nedd4-2 KO was generated in the MDCKI-hNKCC2 cells by CRISPR/Cas9 technology. 3.2 µg sgRNA against the HECT domain of Nedd4-2 (5’-TCTTCCAATAAAAGTGAAAT AGG-3’) (Synthego) and 1 g Cas9 mRNA (TriLink/Tebu-bio) were transfected into MDCKI-hNKCC2 cells using electroporation (4D-NucleofectorTM core unit/X unit, Lonza). 200,000 cells per reaction were pelleted at 300 x g for 5 min, washed twice in PBS pH7.4 and resuspended in OptiMEM media (Gibco). Cells were transferred to the electroporation cuvettes, mixed with Cas9 mRNA and sgRNA and incubated for 1 min before electroporation. Cells were allowed to recover for 3 min at room temperature, before adding 100 µl culture media and transferring to 24 well plastic culture plates. Media was changed after 72 hours and cells were subsequently sub-cultured after 4 days. Genomic DNA was purified from cells (Purelink genomic DNA kit, Thermo Fischer) and PCR performed with primers covering the sequence targeted by the sgRNA (forward primer: 5’-ggcgtttctcatcctcccaa-3’, reverse primer: 5’-ccacaaacagcaatgtccca-3’). The resulting PCR product was sequenced (Eurofins MWG) and analyzed for indels and knockout score using the ICE tool (https://www.editco.bio/crispr-analysis).

#### Semi-quantitative reverse transcriptase-PCR and standard RT-PCR

Semi-quantitative reverse transcriptase-PCR (RT-qPCR) and standard RT-PCR were performed as previously described (44). Primers against *SLC12A1* (forward: 5’-ACTATCGTAACACCGGCAGC-‘3; reverse: 5’-CAGCTGAACTTGGGGTGACT-‘3) and *NEDD4L* (forward: 5’-CACTGGAGGGTGCCAAGGAT-‘3; reverse: 5’-CCGTTGGGCGCTATCCTCAT-‘3) were used. For normalization primers against *18S rRNA* were used (forward: 5’-GGATCCATTGGAGGGCAAGT-‘3; reverse: 5’-ACGAGCTTTTTAACTGCAGCAA-‘3).

#### Immunoblotting and antibodies

Protein samples were separated using 4–15% gradient polyacrylamide gels (Criterion TGX Precast Protein Gels, BioRad) and transferred electrophoretically to PVDF membranes. Immunoblotting was performed using standard methods. Primary antibodies used are; rabbit anti-NKCC2 1495 (45, 46), rabbit anti-NKCC2 L320 (47), rabbit anti-pT96/T101-NKCC2 (48), anti-S126-NKCC2 (19), rabbit anti-Nedd4-2 A27 (49), rabbit anti-FLAG (F7425, Millipore), rabbit anti-proteasome 20s (Ab3325 Abcam), rabbit anti-beta actin (A2066, Sigma) and mouse anti-ubiquitin (P4D1, Cell Signalling).

#### Deglycosylation assay

Cells were grown on plastic until confluent, induced, washed in PBS-CM (PBS containing 1 mM CaCl_2_ and 0.1 mM MgCl_2_, pH 7.5) and harvested in dissection buffer (300 mM sucrose, 25 mM imidazole, 1 mM EDTA, pH 7.2) containing protease inhibitors (5 μg/ml leupeptin, 100 μg/ml Pefabloc). Samples were sonicated and centrifuged for 10 min at 10,000 × g at 4 °C. Protein concentrations were measured using BCA assays (Thermo) and equal protein amounts were denatured in 1× denaturing buffer (New England Biolabs) for 15 min at 65 °C, chilled on ice and centrifuged for 10 sec, before deglycosylation using PNGase F (Peptide N-glycosidase F) or EndoH (Endoglycosidase H) (New England Biolabs) for 1 h using standard protocols. 4× Laemmli gel sample buffer (250 mM Tris base, pH 6.8, 34.8% glycerol, 8% SDS, 4% bromphenolblue with 60mg/ml Dithiothreitol) was added, and samples were heated for 10 min at 65 °C before SDS-PAGE.

#### Immunocytochemistry

Cells were seeded on semi-permeable transwell plates (Corning #3450 or Greiner #657641), grown in complete media until confluent and induced. Cells were fixed in 4% paraformaldehyde/PBS for 15 min at room temperature, before permeabilization in 0.3% SDS in PBS for 5 min. Labeling was performed as described (42) and a Leica TCS SL confocal microscope with an HCX PL Apo 63× oil objective lens (numerical aperture, 1.40) was used for obtaining image stacks with a *z*-distance of 0.1 μm between images.

#### Cell surface biotinylation

Cells were seeded on semi-permeable transwell plates and grown in complete media until confluent. Cells were induced, washed twice in ice-cold PBS-CM and apical membrane proteins were biotinylated with 1.0 mg/ml of sulfosuccinimidyl 2-(biotin-amido)-ethyl-1,3-dithiopropionate (EZ-link Sulfo-NHS-SS-biotin, Invitrogen) in ice-cold biotinylation buffer (10 mM triethanolamine, 2 mM CaCl_2_, 125 mM NaCl, pH 8.9) for 30 min at 4 °C with mild agitation. Cells were washed twice with quenching solution (50 mM Tris-HCl in PBS-CM, pH 8) and once in PBS-CM to remove the excess of biotin, before being lysed in lysis buffer (50 mM Tris-HCl, 150 mM NaCl, 0.1% Triton X-100, 5 mM EDTA (pH 7.5), 20 mM N-ethylmaleimide (Sigma), 22 μM PR619 (Abcam), 5 μg/ml leupeptin, 100 μg/ml Pefabloc, and PhosSTOP phosphatase inhibitor tablets (Roche Diagnostics). Lysates were sonicated and centrifuged at 10,000 × g for 5 min at 4°C. One fraction of the supernatant was retained for total protein estimation (total fraction). The remaining supernatant was incubated for 1 h in spin columns with 200 μl of Neutravidin Plus UltraLink Resin (Invitrogen) under rotation at room temperature. Neutravidin beads were washed in PBS (pH 7.2) with inhibitors and biotinylated proteins eluted with 1x Laemmli sample buffer with a final DTT concentration of 15 mg/ml.

#### Cell surface biotinylation coupled with immunoprecipitation (IP)

Surface biotinylation and immunoprecipitation protocols were followed (34), except biotinylated proteins were eluted using 50 mM DTT in IP lysis buffer for 1 h at RT with rotation before immunoprecipitation with FLAG-M2 affinity beads.

#### NKCC2 protein half-life assays

Cells were grown on semi-permeable transwell plates in complete media until confluent and induced. Cells were washed once with DMEM high glucose and incubated in 355 µM cycloheximide (Sigma) and 5 µM Actinomycin D (Sigma) at 37 °C for various time points. Cells were washed in PBS-CM pH 7.5 and lysed in Laemmli buffer with final concentration of 15 mg/ml DTT. Samples were sonicated and subjected to immunoblotting. For calculation of the protein half-life, average band densities for each time point were normalized to time zero and fitted using nonlinear regression and a one-phase exponential decay equation using GraphPad Prism software.

#### Thallium-ion (Tl+) flux transport assay

NKCC2 activity was accessed using the FluxOR II Green Potassium Ion Channel Assay kit (Thermo Fisher Scientific). Cells were seeded in black 96-well plates with clear bottom (PerkinElmer) and induced. The thallium-ion flux assay was performed as previously described (50). For normalization to total protein concentration, all buffers were removed after the assay, and cells were lysed using 30 µl lysis buffer containing protease and phosphatase inhibitors. Protein concentration was determined using a bicinchoninic acid (BCA) protein assay (Pierce) according to the manufacture protocol.

#### Biotin-based internalization assay

Performed as previously described (34, 42) with the following modifications. Confluent cells grown on filter plates were induced and washed once in ice-cold PBS-CM and once in ice-cold biotinylation buffer (10 mM triethanolamine, 2 mM CaCl_2_, 125 mM NaCl, pH 8.9). To label apical surface NKCC2, the apical compartment of cells was incubated for 30 min at 4 °C in ice-cold biotinylation buffer containing 1 mg/ml EZ-link Sulfo-NHS-SS-biotin (Pierce) with mild agitation. Cells were washed once in ice-cold quenching buffer (PBS-CM, 50 mM Tris-HCl, pH 8) followed by one wash in PBS-CM. At this point NKCC2 surface expression controls were made following the standard biotinylation protocol. The remaining cells were incubated in pure DMEM media for 0, 15 or 30 min at 37 °C to allow plasma membrane proteins to internalize. At each time-point, endocytosis was stopped by rapidly cooling the cells using ice-cold PBS-CM and biotin was stripped from the remaining surface proteins by 3 × 20 min incubations at 4 °C with the membrane-impermeable reducing agent sodium-MES (MesNa, 200 mM) in (100 mM NaCl, 1 mM EDTA, 50 mM Tris, pH 8.6, 0.2% BSA). Reactions were quenched with 120 mM iodoacetic acid in PBS-CM followed by one PBS-CM wash. Cells were lysed and biotinylated proteins isolated according to the normal surface biotinylation protocol.

## Results

### Characterization of tetracycline inducible MDCKI-hNKCC2 cells

After induction with tetracycline, MDCKI-hNKCC2 cells had robust NKCC2 mRNA expression, but NKCC2 protein abundance was relatively weak (Figure 1A). Addition of valproic acid (5 or 15 mM), a histone deacetylase (HDAC) inhibitor (51), for 24 h before tetracycline induction resulted in robust expression of mature and high molecular weight NKCC2, including a potential dimeric form (Figure 1B). A combined tetracycline and 5 mM valproic acid treatment increased levels of mature NKCC2 after 16 h (Figure 1C), and 24 h of dual induction was used for the subsequent studies. No other proteins examined had increased expression with valproic acid (not shown). Cell-surface biotinylation or immunolabeling of MDCKI-hNKCC2 cells grown on filters demonstrated that NKCC2 was, under basal conditions, localized to the apical plasma membrane with additional NKCC2 in intracellular compartments (Figure 1D and 1E).

**Figure 1.**
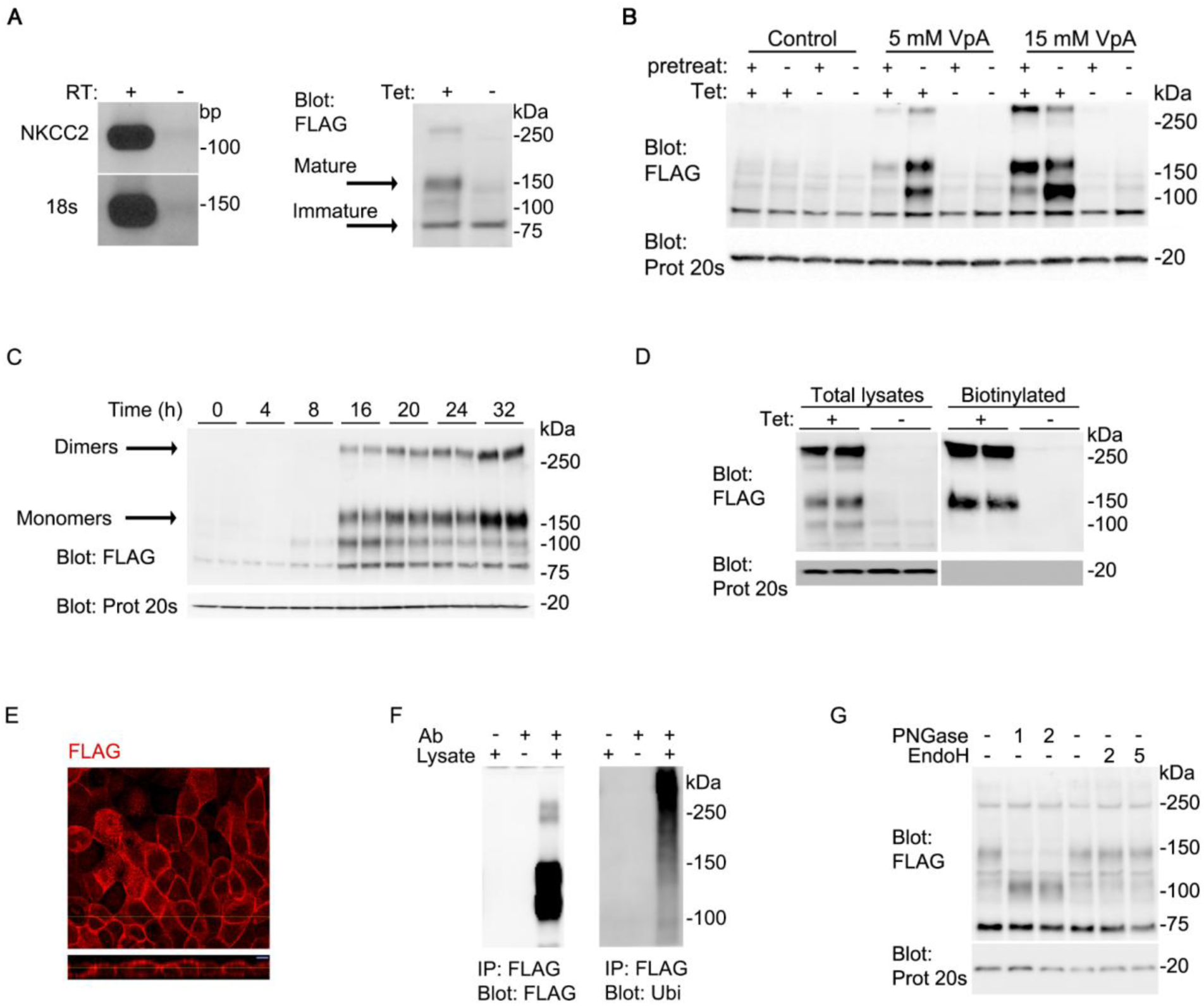
Generation of a tetracycline inducible NKCC2 MDCKI cell line. A) After treatment of MDCKI-hNKCC2 cells with tetracycline, RT-PCR with primers against hNKCC2A and 18S rRNA (control) demonstrated robust NKCC2 mRNA expression. Immunoblotting of MDCKI-hNKCC2 cells using a flag-tag antibody demonstrated relatively low NKCC2 protein levels. B) Representative immunoblots using flag-tag or 20S proteasome (control) antibodies of MDCKI-hNKCC2A cells pretreated with 5 or 15 mM valproic acid (VpA) for 24 h prior to tetracycline (Tet) induction. C) Immunoblots of MDCKI-hNKCC2A cells treated with tetracycline combined with VpA for various time points. D) Representative immunoblots of lysate from tetracycline and VpA induced MDCKI-hNKCC2A cells grown on semi-permeable supports subjected to apical surface biotinylation. Total and biotinylated pools were blotted with flag-tag or 20S proteasome antibodies. E) Representative confocal laser micrograph (top, xy plane; bottom, xz plane) of tetracycline and VpA induced MDCKI-hNKCC2A cells grown on semi-permeable supports labeled with flag-tag antibody. F) NKCC2 was immunoprecipitated (IP) from MDCKI-hNKCC2 cells using a flag-tag antibody, with negative controls performed with or without addition of antibody or lysate. Immunoprecipiates were blotted for flag (NKCC2) or total ubiquitin (Ubi). G) NKCC2 exists as a highly complex glycosylated protein. Representative immunoblots using flag-tag and 20S proteasome (control) antibodies of lysates from tetracycline and VpA induced MDCKI-hNKCC2 cells treated with PNGase F (1 or 2 µl) or EndoH (2 or 5 µl) for 1 h at 37°C.

*In vivo,* the mature function form of NKCC2 is highly glycosylated (52, 53). To assess NKCC2 glycosylation in the MDCKI-hNKCC2 cells, protein lysates were treated with endoglycosidase H (Endo H), which removes high-mannose and hybrid Asn(N)-linked carbohydrates or Peptide:N-glycosidase F (PNGase F), which removes all types of N-linked glycosylation. Endo H had limited effect on NKCC2, whereas PNGase F lowered the observed molecular weight of NKCC2, suggesting that NKCC2 exists as a highly glycosylated protein in the MDCKI-hNKCC2 cells (Figure 1F and 1G).

### Activation of WNK-SPAK/OSR1 or cAMP-mediated pathways increases NKCC2 surface expression

To investigate whether WNK-SPAK/OSR1 or PKA-mediated regulation of NKCC2 is possible in MDCKI-hNKCC2 cells, the cells were treated with hypotonic low chloride solution (activates WNK-SPAK signaling) or forskolin (activates adenylyl cyclase and downstream PKA) and surface NKCC2 levels were accessed by biotinylation. NKCC2 levels on the apical plasma membrane were greater following either treatment (Figure 2A-B). Total and plasma membrane levels of pT96/101-NKCC2 were significantly elevated by stimulation of the WNK-SPAK/OSR1 pathway, whereas forskolin had no significant effect on total or membrane pT96/101-NKCC2 levels (Figure 2C-D). In contrast, forskolin increased pS126-NKCC2 phosphorylation and plasma membrane abundance, whereas hypotonic low chloride had no significant effect (Figure 2E-F). Collectively, these data demonstrate that WNK-SPAK/OSR and cAMP signaling pathways are present in MDCKI-hNKCC2 cells and each pathway can modulate NKCC2 trafficking independently.

**Figure 2.**
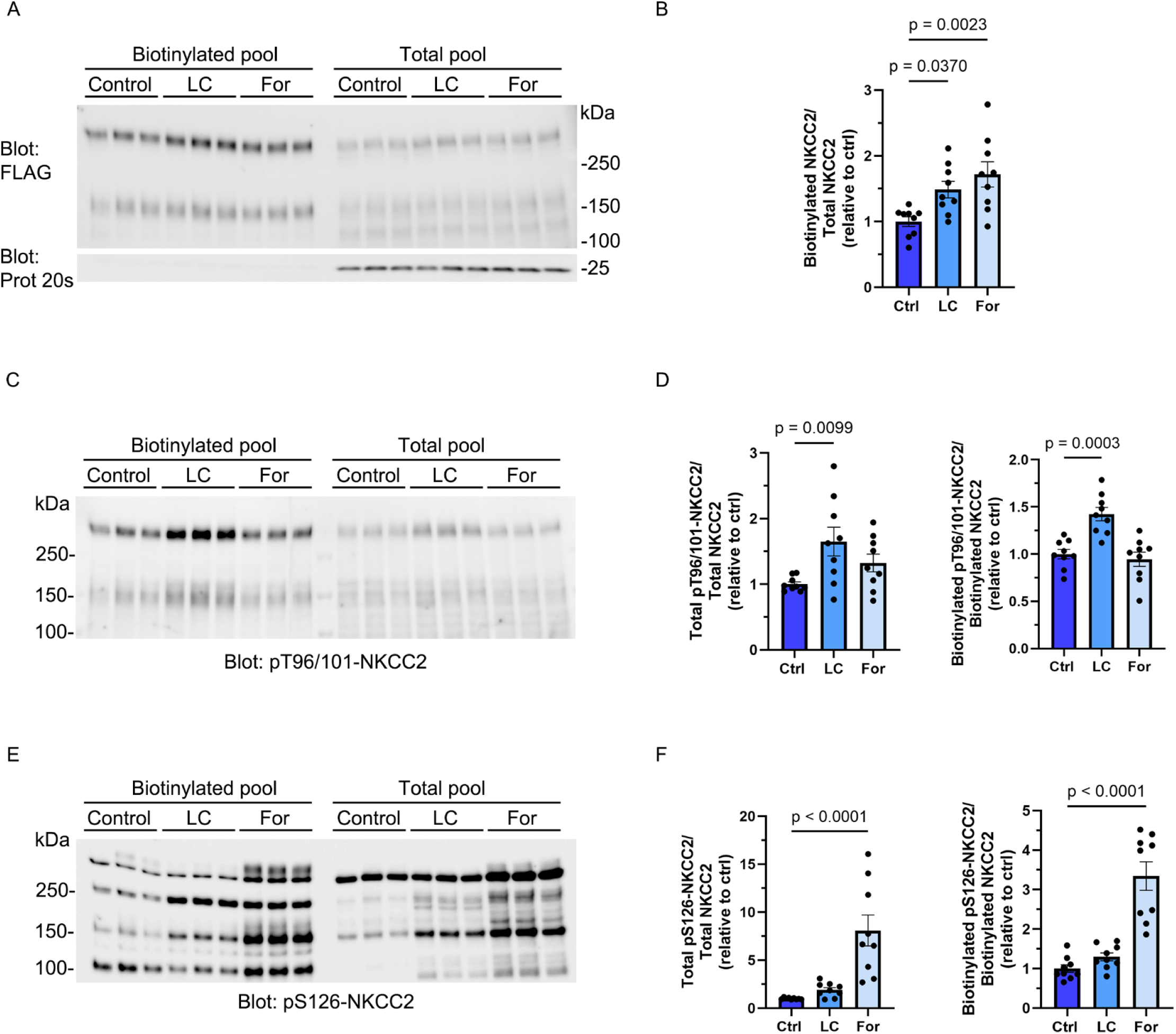
NKCC2 in MDCKI-hNKCC2 cells is regulated by the WNK-SPAK/OSR1 and cAMP pathways. Induced MDCKI-hNKCC2 cells grown on semi-permeable supports and treated with hypotonic low chloride (LC) or forskolin (For) were subjected to apical surface biotinylation. Representative immunoblots of total and biotinylated pools blotted for flag-tag (A), pT96/101-NKCC2 (C), or pS126-NKCC2 (E) with 20S proteasome antibodies (control) are shown. Semi-quantitative assessment of total and apical membrane levels are shown in B, D and E. Data are means ± S.E.M. and analyzed by one-way ANOVA.

### Ubiquitylation at K871 modulates NKCC2 plasma membrane abundance

Mass spectrometry analysis of NKCC2 immunoprecipitated from the MDCKI-hNKCC2 cells identified 15 ubiquitylated lysines. Of these, 5 sites were confirmed to be ubiquitylated in a previous UbiScan (34) and in large-scale mass spectrometry studies of mouse kidney lysates (28) (Supplemental Table 1).

To study the role for site-specific ubiquitylation in NKCC2 regulation, isogenic MDCKI cell lines expressing hNKCC2 deficient in one of the 5 different ubiquitylation sites identified through independent studies (K140, K726, K846, K871, K1012) were generated. The resultant lines had uniform cell morphology, growth characteristics, and robust NKCC2 expression with 10 µg/ml tetracycline and 5 mM valproic acid induction for 24 h (Supplemental Figure 1). Initial cell-surface biotinylation studies demonstrated that NKCC2 was expressed in the apical plasma membrane for all mutants, with K726R and K871R mutants tending to be increased (Supplemental Figure 2A). However, as the K726R mutant showed significantly *lower* levels of total NKCC2 in the MDCKI cell line (Supplemental Figure 2B), suggesting altered biogenesis, only the K871R NKCC2 mutant was analyzed further.

Additional cell-surface biotinylation assays determined that apical plasma membrane NKCC2 levels were ∼50% greater in K871R mutants (Figure 3A). Although significantly reduced NKCC2 ubiquitylation levels at the plasma membrane were also observed in K871R mutants, this had no significant effect on total NKCC2 abundance. RT-qPCR detected no significant differences in constitutive mRNA levels between wt and K871R mutants (Figure 3B).

**Figure 3.**
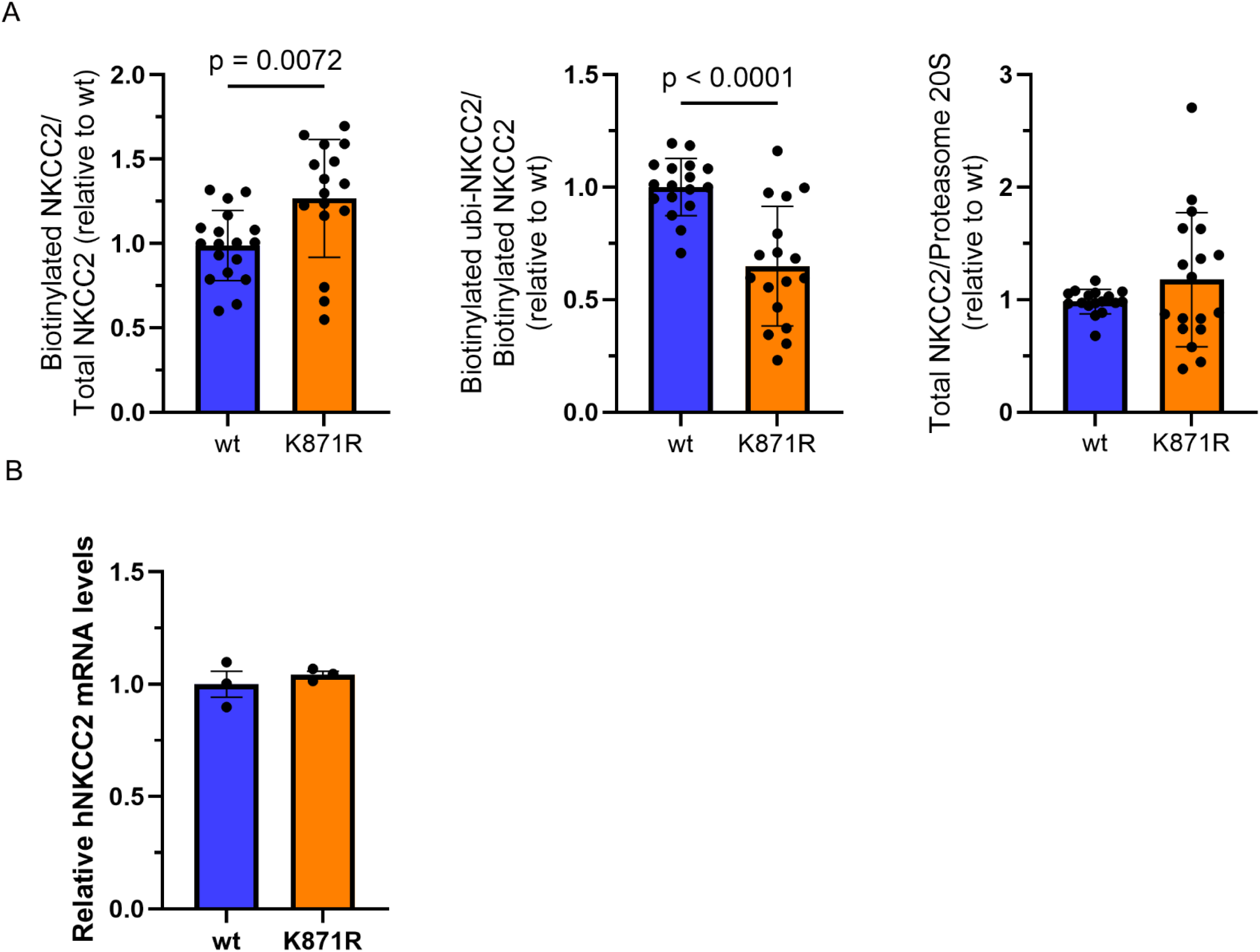
Mutation of the K871 ubiquitylation site increases NKCC2 levels at the plasma membrane. A) Semi-quantitative assessment of apical membrane levels of NKCC2, ubiquitylated NKCC2, and total NKCC2 in wildtype (wt) and K871R cell lines. B) RT-qPCR analysis of mRNA with primers against hNKCC2A and 18S rRNA in wt and K871R cell lines. Data are means ±S.E.M. and analyzed by unpaired Student’s t test.

### MDCKI-hNKCC2-K871R cells have greater thallium uptake

To assess if the greater membrane levels of the K871R mutant relative to wt NKCC2 resulted in higher NKCC2-mediated transport in the cells, we measured thallium (Tl^+^) influx as a surrogate marker for K^+^ uptake (54). To maximize NKCC2 activity via the WNK-SPAK pathway, cells were depleted of intracellular chloride for 1 hour prior to the assay. MDCKI-hNKCC2-K871R cells had a significantly higher Tl^+^ influx relative to MDCKI-hNKCC2 cells. The relative ∼50% increase in TI^+^ uptake in the MDCKI-hNKCC2-K871R cells is in line with the observed fold-increase in NKCC2 plasma membrane levels. When cells were treated with the NKCC2 inhibitor bumetanide, Tl^+^ influxes were significantly reduced to similar levels in both cell lines, indicating that the higher uptake in MDCKI-hNKCC2-K871R cells is driven solely by higher NKCC2-mediated transport (Figure 4).

**Figure 4.**
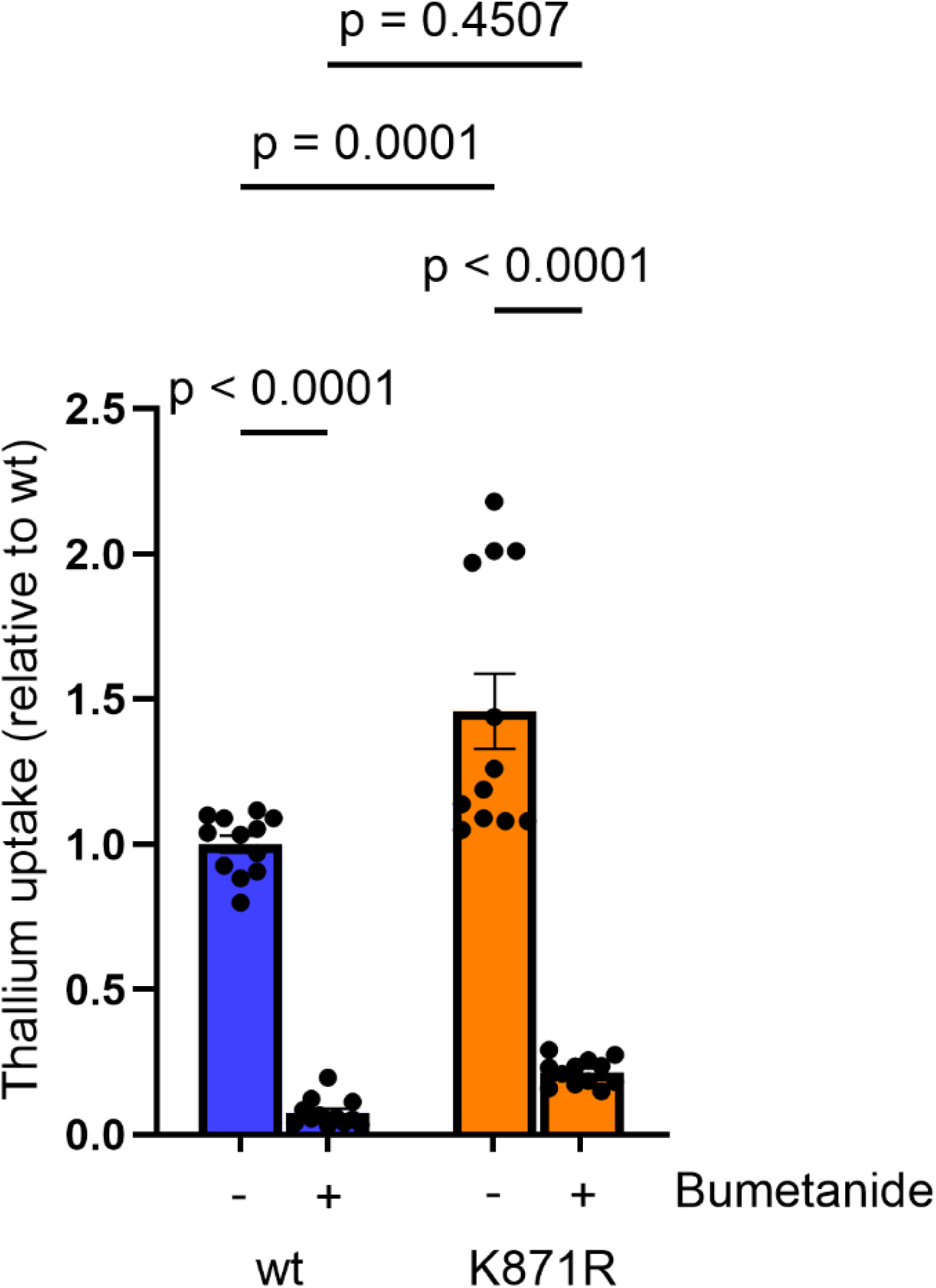
Thallium (Tl+) uptake in wildtype (wt) NKCC2 and K871R-NKCC2. For each individual cell line the uptake (fluorescence intensity) is normalized to the average initial baseline period (without Tl+). Uptakes are normalized to NKCC2 expression per cell line and subsequently normalized to wt NKCC2 without bumetanide treatment. Data are means ± S.E.M, n=12 (3 independent experiments with 4 replicates for each time point per experiment). Data are analyzed by two-way ANOVA followed by a Tukey multiple-comparison test.

### Removing ubiquitylation of K871 does not alter constitutive NKCC2 endocytosis from the apical plasma membrane

Ubiquitylation of membrane proteins can be a stimulus for protein endocytosis (55–57). Furthermore, studies in TAHL suspensions indicate that NKCC2 undergoes constitutive endocytosis and recycling from the plasma membrane (36). We therefore hypothesized that the endocytic rate of NKCC2 would be reduced with elimination of ubiquitylation at K871. NKCC2 internalization assays (42) demonstrated that NKCC2 was constitutively endocytosed from the apical plasma membrane, with ∼15% of NKCC2 internalized after 30 min (Figure 5A). However, in MDCKI-hNKCC2-K871R cells, although the absolute amount of internalized NKCC2 trended lower compared to MDCKI-hNKCC2 cells, this was not significantly different. Furthermore, the rate of NKCC2 internalization did not significantly differ between the lines (Figure 5A).

**Figure 5.**
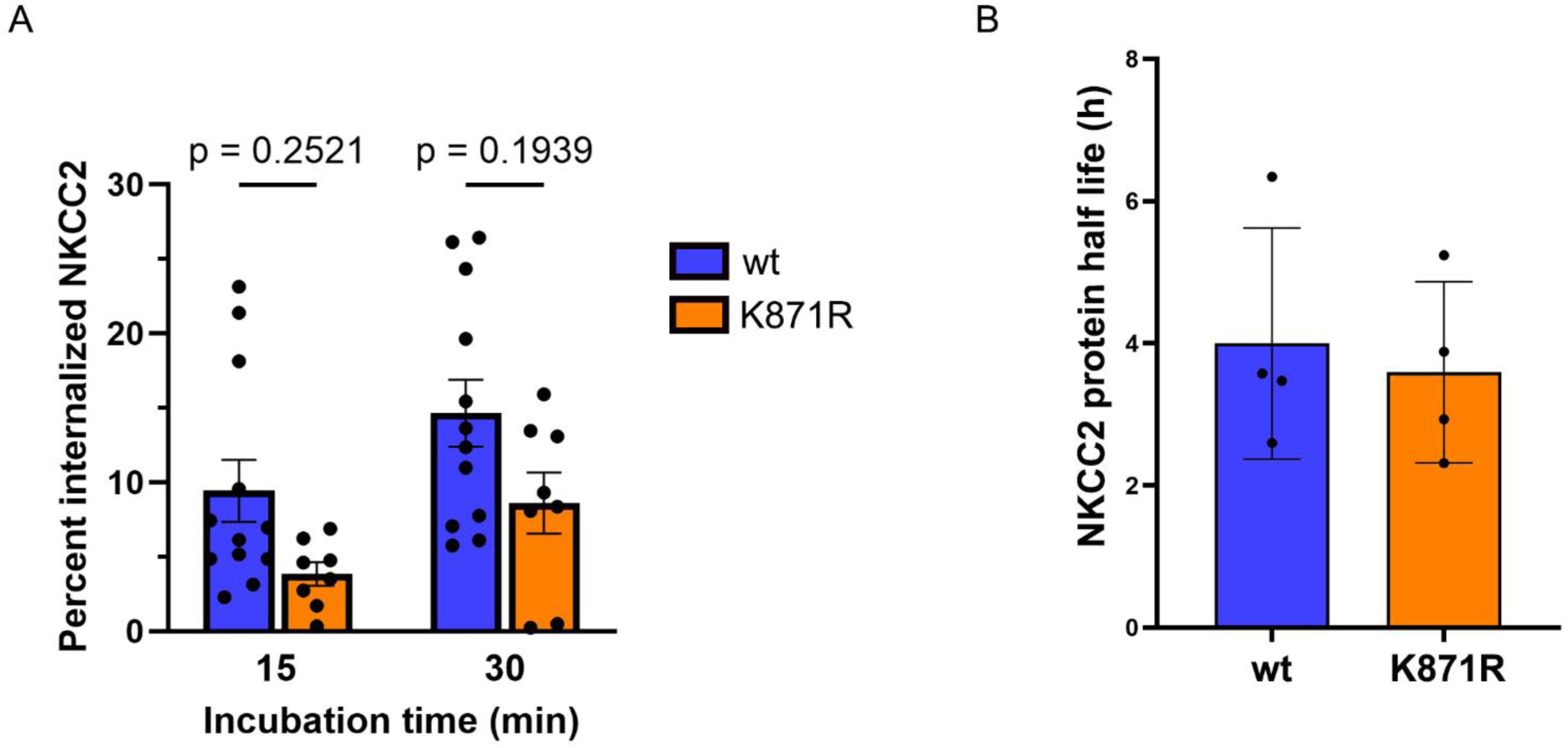
Internalization rates or protein half-life of NKCC2 are not decreased in the K871R mutant cells. A) Semi-quantitative assessment of the percentage of internalized NKCC2 (steady-state surface levels equals 100%) at various time-points. Data are means ±S.E.M (n=9–12) and analyzed using a two-way repeated-measures ANOVA followed by a Holm-Šídák multiple comparison test. P(inter)=0.9158. B) Summary of NKCC2 protein half-life from 4 independent experiments. Data analyzed by unpaired Student’s t-test. Data are means ±S.E.M.

### Ubiquitylation of K871 does not alter NKCC2 degradation

The most widely studied function of protein ubiquitylation is to mark the tagged protein for degradation (55, 58, 59). To assess this role, degradation rates of NKCC2 were analyzed by inhibiting transcription (actinomycin D) and translation (cycloheximide) in MDCKI-hNKCC2 and MDCKI-hNKCC2-K871R cells and assessing NKCC2 levels over time by immunoblotting cell lysates. The average half-life for NKCC2 calculated from 4 independent experiments was 3.6 ± 0.6 h in K871R NKCC2 mutants, which was not significantly different from the 4.0 ± 0.8 h wt NKCC2 half-life (Figure 5B).

### Nedd4-2 interacts with NKCC2 and alters total NKCC2 abundance

Nedd4-2 was co-immunoprecipitated with NKCC2 (Figure 6) confirming previous studies (23). After CRISPR/Cas9 mediated Nedd4-2 deletion in the MDCKI-hNKCC2 cell lines, Nedd4-2 mRNA was significantly reduced by greater than 80%, without any significant effect on NKCC2 mRNA (Figure 7A). At the protein level, Nedd4-2 was undetectable in Nedd4-2 KO cells (Figure 7B). In cells lacking Nedd4-2, cell surface biotinylation studies indicated that plasma membrane NKCC2 abundances were not significantly altered (Figure 8). However, the total levels of NKCC2 were significantly higher after Nedd4-2 deletion.

**Figure 6.**
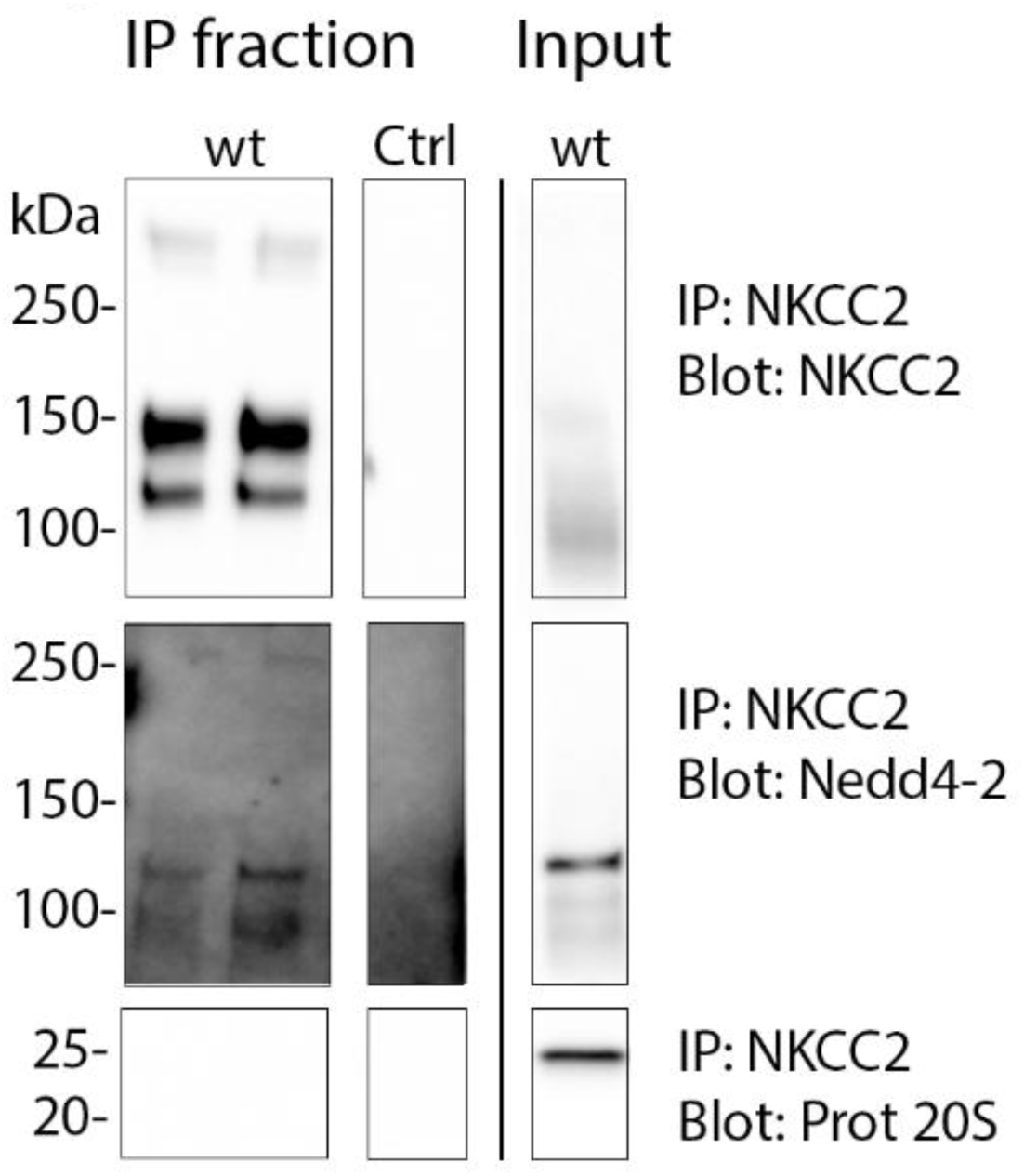
NKCC2 interacts with Nedd4-2 in MDCKI cells. NKCC2 was immunoprecipitated (IP) from MDCKI-hNKCC2 cells (wt) using FLAG-M2 affinity beads and negative controls (Ctrl) were performed in the absence of lysates. Eluate or input fractions were immunoblotted for NKCC2, Nedd4-2 or proteasome 20s (control).

**Figure 7.**
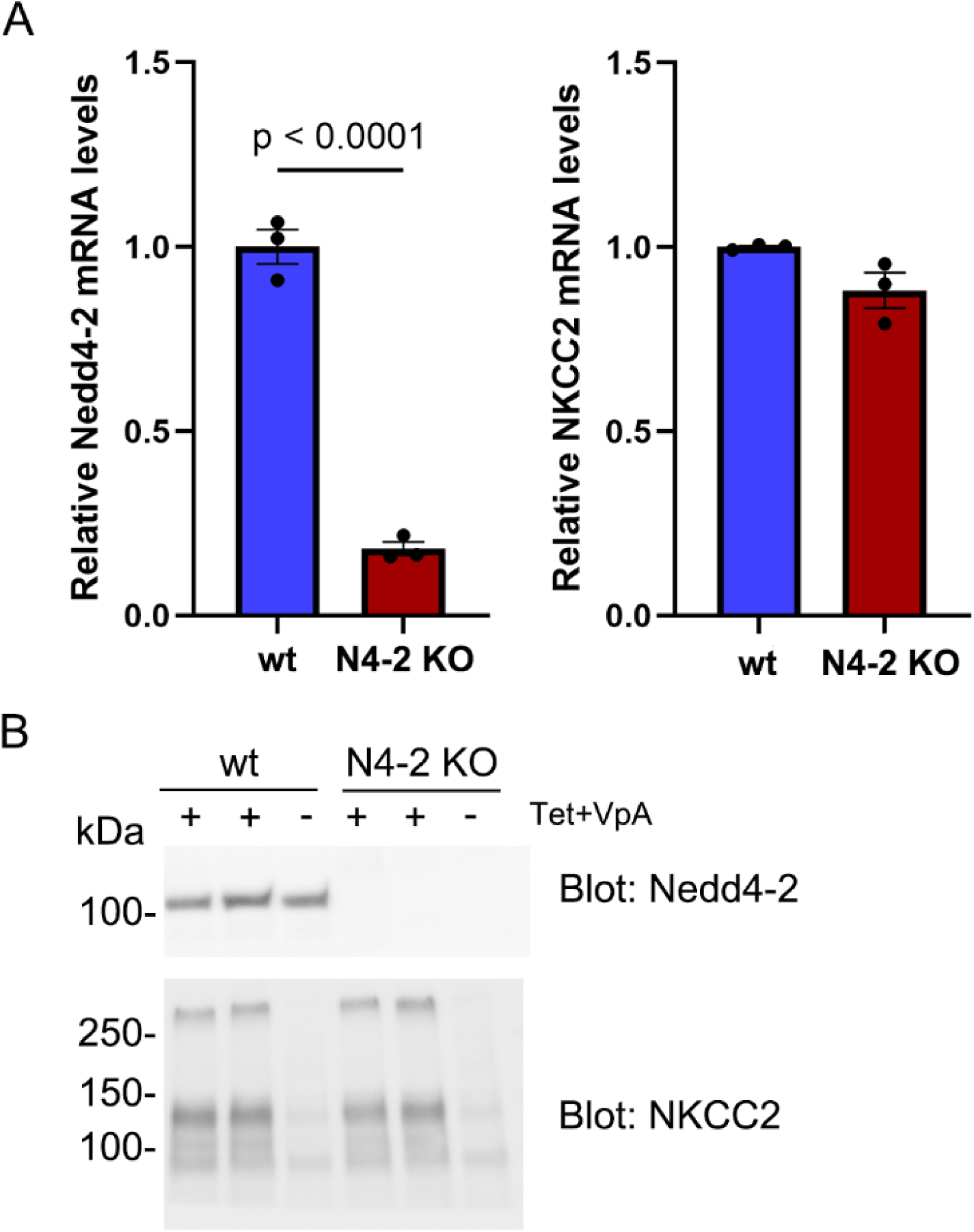
Nedd4-2 mRNA and protein levels in MDCKI-hNKCC2 Nedd4-2 knockout (N4-2 KO) cell lines. A) mRNA levels of Nedd4-2 (left) and NKCC2 (right) in MDCKI-hNKCC2 wildtype (wt) and N4-2 KO cell lines. Data are means ± S.E.M. (n=9) and assessed by unpaired Student’s t test. B) Representative immunoblots of NKCC2 and Nedd4-2 in protein lysates from MDCKI-hNKCC2 wt and N4-2 KO cell lines.

**Figure 8.**
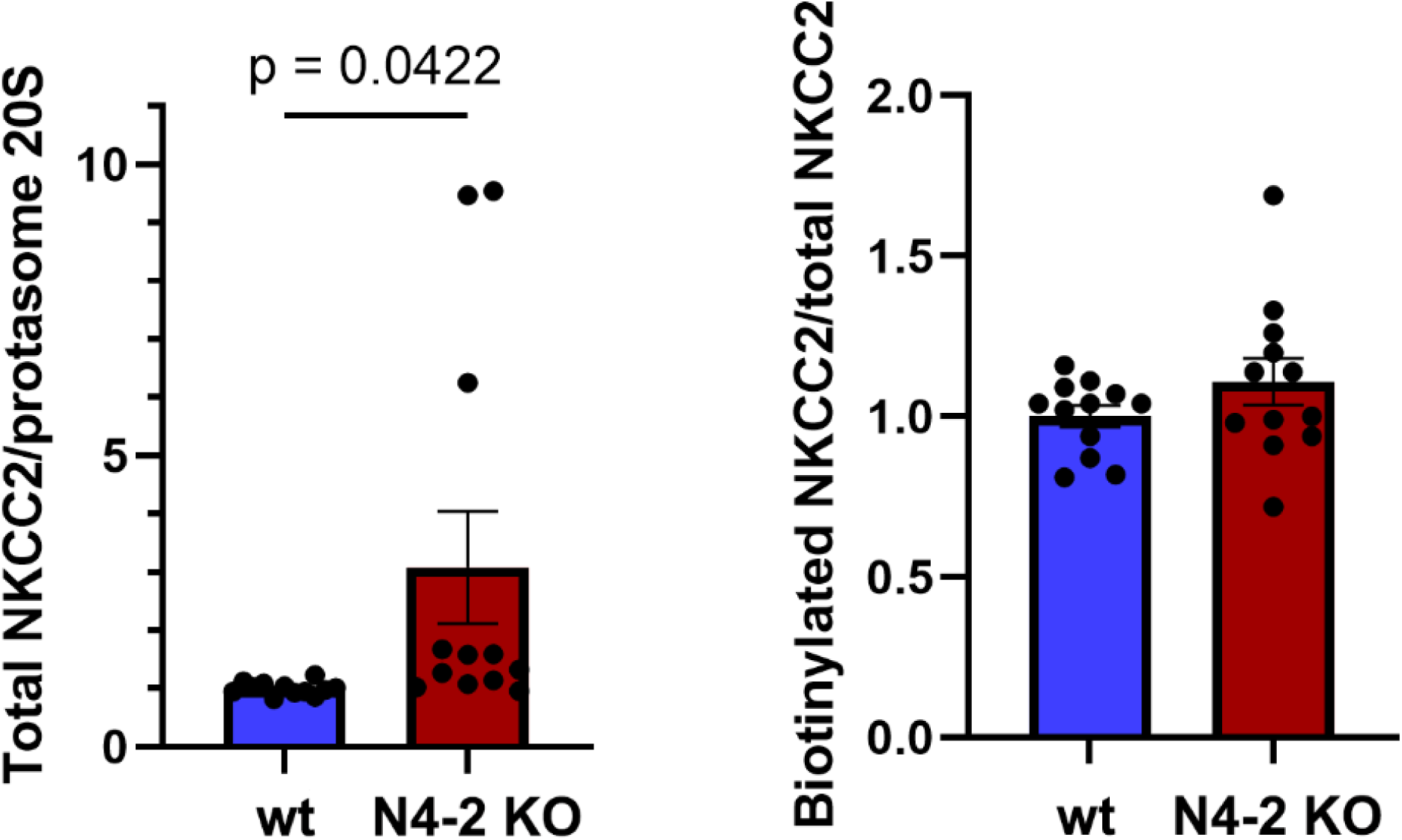
Nedd4-2 knockout increases total NKCC2 levels but not its membrane abundance in MDCKI-hNKCC2 cells. Semi-quantitative assessment of NKCC2 levels in total and plasma membrane fractions of MDCKI-hNKCC2 wildtype (wt) and N4-2 KO cell lines. Data are means ± S.E.M. (n=12) and assessed by unpaired Student’s t test.

## Discussion

Despite its physiological importance (4–6), holes remain in our knowledge of NKCC2 cell biology - particularly that of posttranscriptional regulation - due to a lack of robust mammalian expression systems. As we successfully expressed the sodium chloride cotransporter NCC in MDCK cells using a tetracycline inducible system (33, 34) that minimizes chronic overexpression artifacts and enables precise comparison between site-specific mutants, we hypothesized that such a system was suitable for NKCC2. Furthermore, as the N-terminus of NKCC2 appears to be critical for its expression (38, 39, 60, 61), we hypothesized that addition of an N-terminal Flag-tag would stabilize NKCC2 resulting in robust expression. Therefore, the initial aim of this study was to generate and characterize a tetracycline inducible flag-tagged NKCC2 mammalian cell line suitable for studying NKCC2 biology and allowing direct comparison of mutated forms of NKCC2.

MDCKI-NKCC2 cells had consistent but weak NKCC2 protein induction after tetracycline treatment. However, treatment with the histone deacetylase (HDAC) inhibitor valproic acid (VpA) markedly enhanced NKCC2 expression, consistent with relief of epigenetic repression in the transgene system (51, 62, 63). Notably, valproic acid did not alter the expression of various other endogenous proteins in MDCKI cells, indicating a selective effect on the utilized NKCC2 construct. The underlying mechanism of valproic acid effects remains to be defined, but future studies examining if valproic acid stabilizes NKCC2 mRNA or enhances NKCC2 transcription in a native setting would be informative.

Comparison of previously identified NKCC2 ubiquitylation sites (28, 34) with NKCC2 sites ubiquitylated in the MDCKI-hNKCC2 cell lines suggested that five conserved sites may be important for NKCC2 regulation. Of these, based on initial screens, K871 was selected for extensive investigation and the roles of the other sites currently remain undetermined. Further investigation of the K1012R site is especially warranted, as mutation of this site resulted in an almost doubling of NKCC2 abundance, suggesting a critical role in modulation of NKCC2 stability and degradation.

Ubiquitylation frequently targets proteins to degradation (64). However, no differences were observed in NKCC2 total protein or half-life following K871R mutation. Instead, biotinylation studies demonstrated that elimination of K871 ubiquitylation resulted in greater NKCC2 plasma membrane accumulation, which was supported by elevated thallium uptake in MDCKI-hNKCC2-K871R cells. Whether these effects are direct, due to reduced ubiquitin-dependent NKCC2 endocytosis in the absence of K871 ubiquitylation, or indirect due to effects of the mutation on NKCC2 phosphorylation as seen for other cation chloride cotransporters (34), is currently unknown and requires careful examination of NKCC2 phosphorylation levels in the K871R mutant cells relative to wt cells. Furthermore, the higher plasma membrane abundance in the absence of changes in total protein abundances suggests that NKCC2 is either retained in the plasma membrane or enhanced trafficking of NKCC2 to the membrane occurs in the absence of K871R ubiquitylation. K871R mutants did not have significantly altered membrane internalization rates, suggesting that the first mechanism is unlikely. However, as previous studies have shown that apical exocytosis and recycling of NKCC2 is a regulated process (65), further studies are required to assess if enhanced NKCC2 exocytosis or constitutive recycling occurs in the absence of K871 ubiquitylation.

Although general NKCC2 ubiquitylation is reduced in the K871R mutant, it remains unclear whether this effect is specifically due to the loss of ubiquitylation at K871 or whether the mutation alters the susceptibility of NKCC2 to ubiquitylation at other sites. Appointing site-specific ubiquitylation occupancy is challenging due to limitations in current methods and the possibility of considerable variability in modifying individual sites within the same protein (26). Current LC–MS/MS strategies used to map ubiquitylation sites lack the resolution to determine whether multiple lysine residues on a single NKCC2 protein are modified at the same time, or whether distinct sites are targeted in a location-dependent manner and vary with subcellular compartmentalization. Although the current immunoprecipitation data suggests that multiple sites in NKCC2 are targeted at the same time, as K871R mutation did not completely abolish NKCC2 ubiquitylation, mutating several sites simultaneously has indirect effects on protein function making further investigations of this concept difficult (34).

The E3 ligase Nedd4-2 co-immunoprecipitates with NKCC2 (23). We confirmed this interaction in MDCKI-hNKCC2 cells, and subsequently demonstrated that deleting Nedd4-2 increased cellular NKCC2 protein levels without alterations in its membrane abundance. These findings indicate that Nedd4-2 regulates NKCC2 abundance at the level of protein stability, but does not control its membrane trafficking, suggesting that different E3 ligases mediate site-specific ubiquitylation events. Which sites in NKCC2 are ubiquitylated by Nedd4-2 is unknown and requires further extensive investigation. Whether adaptor and/or scaffold proteins such as 14-3-3 (66, 67) or Ndfip1/2 (68–71) are involved in the Nedd4-2 mediated degradation of NKCC2 has not been investigated.

In conclusion, we have generated a highly valuable inducible polarized kidney cell line model with robust and functional full-length NKCC2 expression. We demonstrate that NKCC2 is highly ubiquitylated and that K871 ubiquitylation plays a role in modulating NKCC2 membrane abundance. Eliminating the E3 ligase Nedd4-2 alters NKCC2 protein stability but not plasma membrane localization. Together, these data support a model in which NKCC2 is regulated by distinct ubiquitylation pathways that independently control protein abundance and membrane trafficking.

## Acknowledgements

We thank Inger Merete S. Paulsen, Helle Høyer, Tina Drejer, Golshah Ayoubi, Cassandra Bennetzen, and Christian V. Westberg for technical assistance. Associate Professor Rasmus O. Bak is thanked for help with CRISPR/Cas9 experiments.

## Funding support

Novo Nordisk Foundation (NNF21OC0067647, NNF20OC0063837 and NNF24OC0095846 to RAF).

Danish Council for Independent Research (3101-00136B to RAF and DFF-4092-00128 to LKR).

Leducq Transatlantic Network of Excellence (17CVD05 to RAF).

## Author contribution

OS, RAF and LKR conceived the study. LNO, AT, QW and LKR conducted experiments and acquired data. LNO, QW, OS, RAF and LKR analyzed and interpreted data. LKR and RAF drafted the manuscript. All authors approved the final version of the manuscript.

## AI statement

Portions of the original draft of the manuscript text were edited with the assistance of generative AI (ChatGPT, OpenAI), such as improving clarity and grammar. No AI tools contributed to study design, data analysis, or interpretation. All final texts were reviewed and approved by the authors.

