## Supplemental material for "Regulation of the Na-K-2Cl cotransporter NKCC2 by ubiquitylation"

**Supplemental Table 1.**

Ubiquitylated lysine residues (K-Ub) on NKCC2 found in MDCKI-hNKCC2 cells compared to sites found in a UbiScan and in large-scale mass spectrometry studies of mouse kidney lysates.

| Protein NKCC2 | MOUSE Position | HUMAN Position | Found in MDCKI-hNKCC2 cells | Found in Rosenbaek and Rizzo et al., 2017 (1) | Found in Wagner et al., 2012 (2) |
|---|---|---|---|---|---|
| **K-Ub** | - | 56 | + | - | - |
| **K-Ub** | 136 | 140 | + | + | + |
| **K-Ub** | 163 | 167 | + | + | - |
| **K-Ub** | 525 | 529 | + | - | + |
| **K-Ub** | 722 | 726 | + | + | + |
| **K-Ub** | 842 | 846 | + | + | + |
| **K-Ub** | 862 | 866 | + | - | + |
| **K-Ub** | 867 | 871 | + | + | + |
| **K-Ub** | 872 | 876 | + | - | + |
| **K-Ub** | 890 | 894 | + | - | + |
| **K-Ub** | 944 | 948 | + | - | + |
| **K-Ub** | 981 | 985 | + | - | - |
| **K-Ub** | 985 | 989 | + | - | - |
| **K-Ub** | 1008 | 1012 | + | + | + |
| **K-Ub** | 1026 | 1030 | + | - | + |

Conserved NKCC2 K-Ubiquitylation sites found in all three studies


1. Rosenbaek, L. L., Rizzo, F., Wu, Q., Rojas-Vega, L., Gamba, G., MacAulay, N. *et al.* (2017) The thiazide sensitive sodium chloride co-transporter NCC is modulated by site-specific ubiquitylation Sci Rep **7**, 12981 10.1038/s41598-017-12819-0
2. Wagner, S. A., Beli, P., Weinert, B. T., Scholz, C., Kelstrup, C. D., Young, C. *et al.* (2012) Proteomic analyses reveal divergent ubiquitylation site patterns in murine tissues Molecular & cellular proteomics : MCP **11**, 1578-1585 10.1074/mcp.M112.017905

**Supplemental Figure 1.**

Immunoblotting of protein homogenates from MDCKI cells expressing NKCC2 K140R, K726R, K846R, K871R, and K1012R mutations. Cells were induced with tetracycline and valproic acid for 24 hours prior to lysis. Protein amounts between mutants are not comparable as they are not run on the same SDS-PAGE gel.

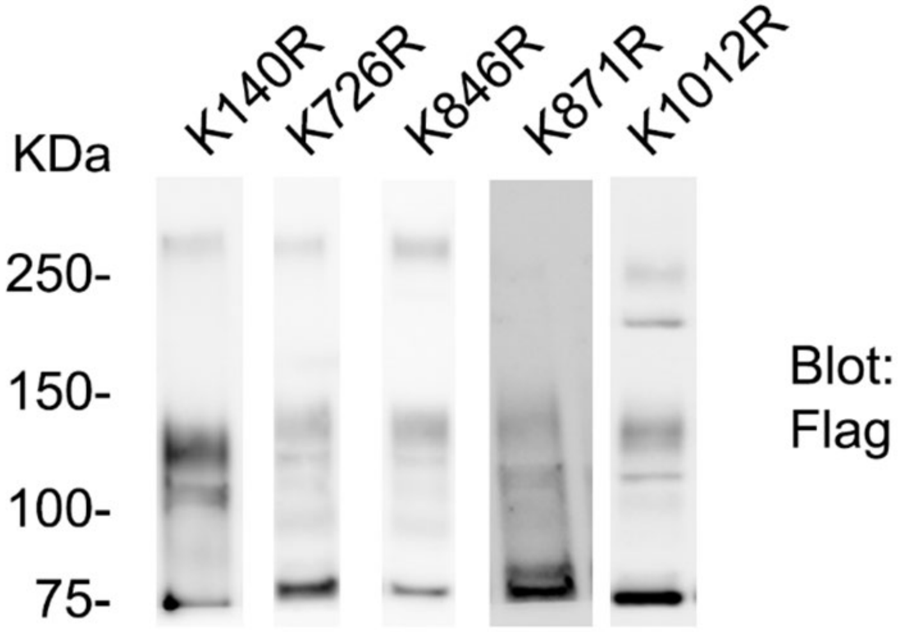

**Supplemental Figure 2.**

Semi-quantitative assessment of apical plasma membrane and total cellular membrane levels in wildtype (wt), K140R, K726R, K846R, K871R, and K1012R mutant NKCC2 cells. Data are means ±S.E.M. (n=4-22) and assessed by ordinary one-way ANOVA with Tukeys post-hoc test.

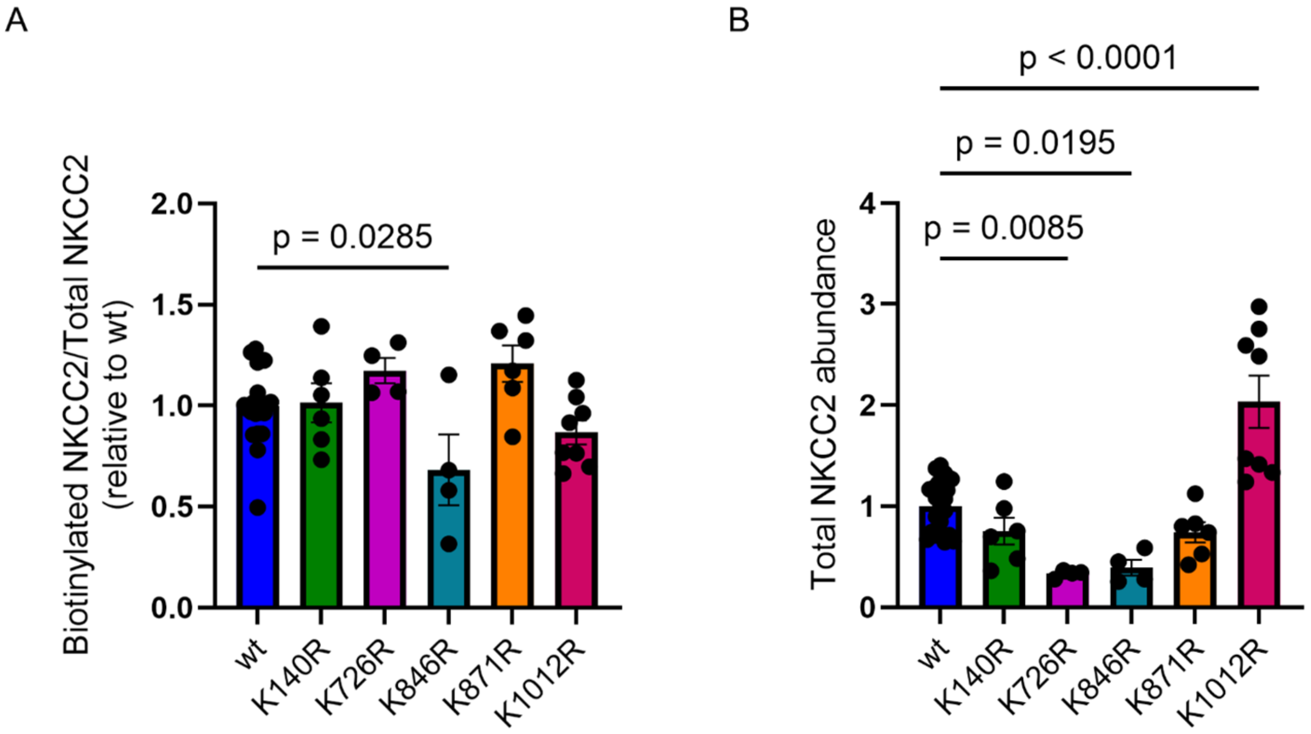
